# Molecular insights into protein disulfide isomerase antagonism by punicalagin

**DOI:** 10.64898/2026.04.29.721714

**Authors:** Osamede C. Owegie, Quinn P. Kennedy, Ivan Hancco Zirena, Oren Levy, Pavel Davizon-Castillo, Moua Yang

**Affiliations:** Bloodworks Northwest Research Institute, Seattle, WA 98102, USA; Centro de Investigación en Medicina de Altura (CIMA), Facultad de Medicina Humana, Universidad de San Martín de Porres, Lima 15001, Peru; Department of Anesthesia, Brigham and Women’s Hospital and Harvard Medical School, Boston, MA 02115, USA; Ben Towne Center for Childhood Cancer Research and the Department of Pediatrics, Seattle Children’s Hospital, University of Washington, Seattle, WA 98102, USA; Division of Hematology and Oncology, Department of Medicine, University of Washington School of Medicine, Seattle, WA 98102, USA

**Keywords:** Thiol isomerase, protein disulfide isomerase, reductase, oxidase, punicalagin, galloylated polyphenol

## Abstract

Punicalagin, an ellagic acid polyphenol from pomegranate, has been proposed as an antagonist of protein disulfide isomerase (PDI) and endoplasmic reticulum resident protein 57 (ERp57), thiol oxidoreductases that regulate protein folding and extracellular thrombotic signaling. Here, biochemical oxidase and reductase assays on PDI show that punicalagin inhibits both activities with micromolar potency, thereby extending earlier work that described only disulfide reductase inhibition. In parallel, thiol labeling of catalytic cysteines revealed no change in the redox state, supporting a noncovalent, allosteric of inhibition. Molecular docking and molecular dynamics simulations showed that punicalagin binds stably and preferentially to defined sites on the N⍰terminal domains of PDI through extensive hydrogen bonding and van der Waals contacts, which is an alternative binding mode to previously reported C-terminal binding. Finally, artificial intelligence-driven network analysis identified PDI as a high-confidence target of punicalagin and related galloylated polyphenols, alongside additional signaling proteins. Together, these findings provide further mechanistic framework for punicalagin-mediated antagonism of PDI and highlight galloylated polyphenols as promising scaffolds for protein disulfide isomerase-targeted therapeutics.

**Highlights:**

- Punicalagin, a galloylated polyphenol, antagonizes not only the reductase activity but also the oxidase activity of protein disulfide isomerase
- Protein disulfide isomerase inhibition by punicalagin is through N-terminal binding
- Punicalagin inhibits conformationally rather than catalytic cysteine modification
- Artificial intelligence network analysis reveals pathway inhibition by punicalagin

## 1. Introduction

Protein disulfide isomerases (PDIs) are multifunctional oxidoreductases that catalyze the formation, reduction, and rearrangement of disulfide bonds in substrate proteins in the endoplasmic reticulum (ER) [1–5]. Structurally, PDIs adopt a modular architecture consisting of four thioredoxin-like domains (**a**, **b**, **b**′, and **a**′) organized in a flexible U-shaped arrangement [6, 7]. The **a** and **a**′ domains each house a CXXC catalytic site, whereas the **b**′ domain forms a hydrophobic substrate-binding cleft that engages unfolded polypeptides, and the c⍰terminal domain stabilizes intramolecular flexibility critical for substrate recognition and folding [7–11]. The thioredoxin-like CXXC active-site motif allows reversible oxidation and reduction of cysteine residues during protein folding [9, 11, 12]. This catalytic plasticity highlights their central role in maintaining cellular proteostasis, ensuring that nascent or stress-denatured polypeptides achieve correct disulfide pairing and structural integrity.

Beyond their classical ER function, PDIs also function extracellularly and at the cell surface, where they modulate redox-dependent receptor activity, integrin function, platelet adhesion, and coagulation thereby contributing to thrombus formation and vascular signaling [13–18]. The discovery of extracellular PDI activity has expanded its functional repertoire from an intracellular folding catalyst to a pivotal regulator of redox communication in vascular physiology [19–21]. As such, pharmacological inhibition of PDI has emerged as a therapeutic strategy in limiting thrombus formation. Over the past decade, efforts to inhibit PDI have yielded a broad spectrum of small-molecule and natural product inhibitors. Among naturally occurring inhibitors, galloylated polyphenols have attracted particular attention due to their distinctive redox-reactive architecture. Comprising one or more galloyl (trihydroxybenzoyl) groups, these compounds exhibit high electron density that enables them to scavenge reactive oxygen species, chelate metal ions, and interact with thiol-containing protein targets [22–27]. These polyphenols combine antioxidant protection with the ability to modulate enzymatic redox activity, positioning them as natural scaffolds for enzyme regulation [23, 25, 27, 28].

Punicalagin, an ellagitannin abundantly present in pomegranate peel (*Punica granatum*), Indian almond leaf extracts (*Terminalia catappa*), and valvet bushwillow bark (*Combretum mole*), stands out as a structurally complex and bioactive galloylated polyphenol with robust antioxidant and anti-inflammatory properties [26, 28–33]. Punicalagin contains a core ellagic acid connected by galloylated motifs to a sugar (**Figure 1A**). We previously found that punicalagin are part of several classes of galloylated polyphenols that have broad activity against thiol isomerases and are antithrombotic when tested in mouse models of arterial thrombus formation [27, 28]. Prior biochemical and computational studies have suggested punicalagin as a noncovalent inhibitor of PDI, demonstrating micromolar binding, inhibition of disulfide reductase activity, and ligand-induced alterations in protein thermal stability and conformational behavior. [34, 35] In particular, comparative analyses have localized punicalagin binding to redox-active domains [34, 35] Yet, the molecular basis of PDI inhibition, its effects on PDI’s oxidase activity or how it affects the redox state of its catalytic cysteines remains incompletely understood. A clearer understanding of this interaction could provide insight into PDI redox regulation and guide the rational design of natural product–based modulators for vascular and oxidative pathologies.

**Figure 1.**
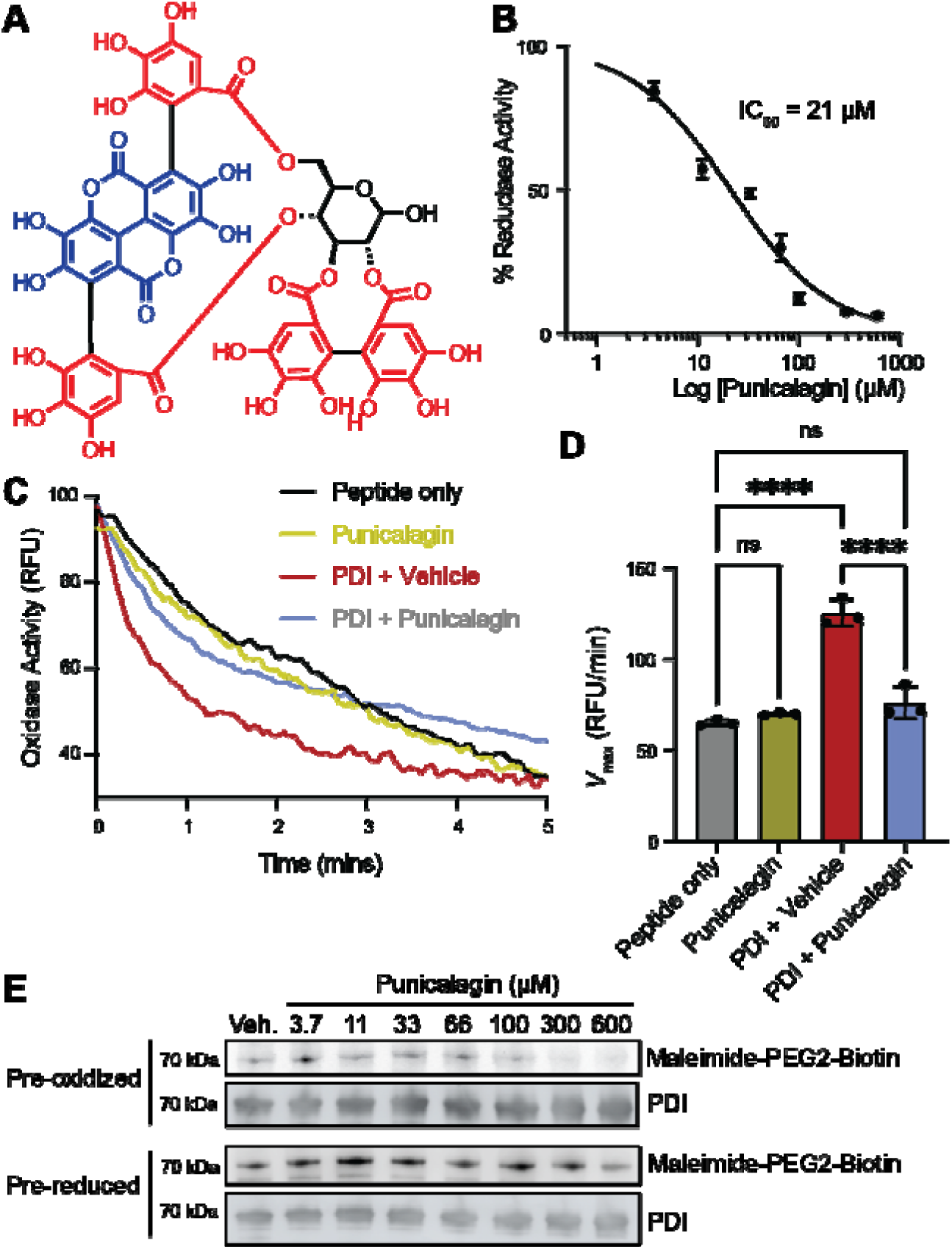
Galloylated polyphenol punicalagin inhibits the oxidoreductase activity of PDI independent of the catalytic cysteines. (A) Structure of punicalagin showing the core ellagic acid (blue), gallic acids (red), and sugar moiety (black). (B) IC_50_ profiling of punicalagin inhibiting the insulin reductase activity of PDI. Recombinant PDI was incubated with the indicated concentrations of punicalagin for 15 minutes prior to reductase activity monitoring by evaluating the ability of PDI to cleave disulfides in insulin. Insulin precipitation was monitored over time at 650 nm absorbance. Data represented as mean ± SEM of 3 replicates. (C) Punicalagin inhibits PDI oxidation of a synthetic NRCSQGSCWN 10-mer peptide. Recombinant PDI was incubated with 40 µM punicalagin in a redox buffer (2 mM GSSG:8 mM GSH; 1:4 molar ratio) prior to the addition of NRCSQGSCWN. Tryptophan fluorescence quenching was monitored over time at ex/em: 280/350 nm. (D) Maximal velocity (Vmax) of fluorescence quenching was quantified from panel (C). (E) Punicalagin does not reduce or oxidize the catalytic CGHC motif of PDI. 10 µM PDI was pre-reduced or pre-oxidized with 200 µM TCEP or H_2_O_2_, respectively, prior to desalting to eliminate excess reductants or oxidants. The protein was then incubated with the indicated concentrations of punicalagin for 30 min. Maleimide-PEG-biotin was then added to quench the reaction and label the free thiols. Biotin and PDI were detected by western blot.

In this study, we investigated the inhibitory mechanism of punicalagin on human PDI using a combined biochemical, computational, and artificial intelligence–based analysis. We employed oxidase and reductase assays in conjunction with thiol labelling methods to determine the redox potency of punicalagin and its inhibitory capacity on PDI. Molecular docking and extensive molecular dynamics simulations were employed on the whole protein to characterize binding interfaces and conformational transitions upon ligand engagement. Finally, we analyzed protein-ligand interaction networks using integrative AI mapping (Plex) [36] to contextualize mechanistic patterns within broader redox and structural landscapes. Together, these approaches aim to elucidate how punicalagin modulates PDI’s catalytic function and to establish a foundation for exploiting galloylated polyphenols as natural allosteric inhibitors in redox-centered therapeutic development to prevent thrombosis.

## 2. Materials and methods

### 2.1. PDI Expression and Purification

The gene encoding human PDI was cloned between the NdeI and BamHI restriction sites in the pT7-FLAG-SBP-1 expression vector (Sigma-Aldrich), which encodes an N-terminal FLAG tag and a C-terminal Streptavidin Binding Protein (SBP) tag. The recombinant plasmid was transformed into Escherichia coli BL21(DE3) (New England Biolabs) and cultured in terrific broth (TB) supplemented with ampicillin, potassium phosphate monobasic, and potassium phosphate dibasic at 37°C. Protein expression was induced overnight at 23°C with 0.5 mM isopropyl β-D-1-thiogalactopyranoside (IPTG). Cells were collected by centrifugation, resuspended in lysis buffer, and lysed by sonication. The lysate was clarified by centrifugation and the supernatant was incubated with Pierce High-Capacity Streptavidin Agarose (Thermo Scientific) for affinity capture of the SBP-tagged PDI. Elution was performed with biotin-containing buffer; eluates were subsequently desalted into phosphate-buffered saline (PBS) to remove biotin. Purified protein was stored at –80°C.

### 2.2. PDI Oxidase Activity Assay

Oxidase activity assays were performed in buffer containing 100 mM KH₂PO₄ and 2 mM EDTA (pH 7.0). A 5 mM stock solution of peptide substrate was prepared in 30% acetonitrile, 0.1% trifluoroacetic acid (TFA), and deionized water to maintain the peptide in a reduced state and minimize spontaneous oxidation. Reduced (GSH) and oxidized (GSSG) glutathione were each prepared as 100 mM stocks in assay buffer. Punicalagin (Sigma) was first dissolved in deionized water to a concentration of 10 mg/mL (92 mM), and the working stock was further diluted to 9.2 mM with assay buffer. Assay buffer was first added to a 3 mL fluorescence cuvette, followed by 240 µL of 8 mM GSH, 80 µL of 2 mM GSSG, recombinant PDI (final concentration 0.8 µM; volume adjusted according to stock concentration), and 13.4 µL of the punicalagin stock (9.2 mM) to achieve a final concentration of 40 µM in punicalagin-treated samples.

Fluorescence measurements were conducted in kinetic mode (excitation 280 nm, emission 350 nm) using a Cary Eclipse Fluorimeter (Agilent). After blanking, the reaction was initiated by adding 8.16 µL of the peptide substrate to a final concentration of 13.6 µM. Fluorescence was recorded immediately to monitor reaction kinetics.

### 2.3. PDI Reductase Activity Assay

Reductase activity assays were performed in buffer containing 61 mM K₂HPO₄, 39 mM KH₂PO₄, and 2 mM EDTA (pH 7.0). For inhibition studies, recombinant PDI was diluted in assay buffer to 900 nM (final concentration: 450 nM per well). Dithiothreitol (DTT) was freshly prepared as a 100 mM stock and diluted in assay buffer to 1.6 mM (final: 0.4 mM per well). The insulin stock (1.5 mM) was diluted to 0.4 mM (final: 0.1 mM per well). Punicalagin was prepared as 10× stocks in assay buffer to yield final concentrations of 0, 3.7, 11, 33, 66, 100, 300, and 600 μM. For each condition, 100 μL of PDI (or buffer for blanks) and the appropriate volume of punicalagin or vehicle were combined in triplicate wells of a clear 96-well plate. The volume of DTT buffer was adjusted per well to maintain a constant final DTT concentration.

Reactions were initiated by adding 50 μL of 0.4 mM insulin to each well. Absorbance at 650 nm was measured at one-minute intervals for up to two hours using a SpectraMax iD5e plate reader (Molecular Devices) to monitor insulin precipitation. IC₅₀ values were derived from plots of normalized PDI activity versus inhibitor concentration and calculated by non-linear regression using GraphPad Prism v10 (La Jolla, CA, USA).

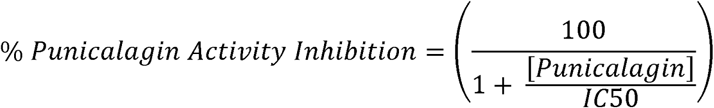

### 2.4. Free Thiol Detection

Free thiols on recombinant human PDI were quantified using a maleimide labeling assay. Recombinant PDI was diluted in PBS (pH 7.4) to 10 µM. For pre-oxidized samples, PDI was incubated with 100 µM H₂O₂ for 15 min at room temperature and then desalted into fresh PBS. For prereduced samples, PDI was incubated with 100 µM TCEP for 15 min at room temperature and desalted similarly. Punicalagin was prepared as 10× stocks in PBS to yield final concentrations of 0, 3.7, 11, 33, 66, 100, 300, and 600 µM. Pre-oxidized or pre-reduced PDI was then incubated with each concentration of punicalagin for 30 min at 37°C. Maleimide–PEG2–biotin was added to a final concentration of 400 µM, and samples were incubated for 15 min at room temperature in the dark. Reactions were quenched with reducing SDS–PAGE sample buffer containing β-mercaptoethanol. Labeled proteins were resolved by reducing SDS–PAGE, transferred to nitrocellulose membranes, and probed with streptavidin–HRP to detect biotinylated thiols; total PDI was assessed in parallel by anti-PDI immunoblotting. Chemiluminescent signals were imaged and quantified using a LI-COR imager (Model M).

### 2.5. Molecular Docking

For unbiased (blind) docking, the entire structure of protein disulfide isomerase (PDI) was enclosed within a cubic grid box of 120 × 120 × 120 Å, providing comprehensive structural coverage. Grid parameter files were generated to define the search space and docking conditions. Docking simulations were performed using the AutoDock Vina algorithm, which evaluates binding affinities and ligand conformations with a scoring function. A total of nine docking poses were generated and subsequently ranked based on predicted binding energies.

To determine the most probable interaction sites, energies and binding orientations from the docking runs were analyzed systematically. Protein–ligand interaction profiling was conducted in Discovery Studio Visualizer, allowing mapping of key residue contacts and hydrogen-bonding patterns. This detailed analysis provided insight into potential binding regions critical for punicalagin–PDI interactions.

### 2.6. Molecular Dynamics Simulation

The best-scoring PDI–punicalagin complex from docking was used as the starting structure for MD simulations. Ligand parameterization for GROMACS topology was performed using AmberTools and ACPYPE. The protein–ligand complex was solvated in a triclinic box using an SPC water model, with at least 10 Å between the solute and box edge to prevent artifacts. Counterions (Na+ and Cl–) were added for system neutralization and to reach a physiological salt concentration (0.15 M NaCl) using GROMACS’ genion tool. CHARMM36 force field was applied for proteins and the system.

Energy minimization was conducted with the steepest descent algorithm over 5000 steps or until the maximum force dropped below 1000 kJ/mol/nm. The system was equilibrated first under NVT conditions, gradually heating from 0 to 300 K over 100 ps with position restraints, followed by an NPT equilibration for 100 ps with pressure maintained at 1 bar using the Parrinello-Rahman barostat and gentle release of position restraints. Production runs were performed for 500 ns with a 2 fs integration timestep, saving trajectory snapshots every 10 ps for analysis.

MD analyses included calculation of root-mean-square deviation (RMSD), root-mean-square fluctuation (RMSF), and persistent hydrogen bonds between PDI and punicalagin using GROMACS utilities. The stability and conformational changes of the complex were visualized with Discovery Studio Visualizer.

### 2.7. Plex Artificial Intelligence (AI)

Punicalagin (PC) and PGHG were introduced as search queries into Plex AI platform (https://www.plexresearch.com) for a chemical structure similarity analysis (parameters: similarity threshold - 0.75, fingerprint type – sim, measure – Tanimoto). Plex results yielded focal graphs, which were exported and imaged via Gephi (https://gephi.org).

### 2.8. Statistical Analysis

All statistical comparisons and data analysis were performed using GraphPad Prism 10. For quantitative assays and kinetic endpoints, experiments were conducted in at least triplicate, and results were presented as mean ± SEM. Dose–response relationships and inhibition statistics were evaluated by one-way ANOVA or non-linear regression, as appropriate. A p-value of < .05 was considered statistically significant.

## 3. Results

### 3.1. Punicalagin inhibits PDI’s oxidoreductase activity without altering the redox state of its active-site cysteines

We previously reported that punicalagin inhibits the ability of PDI to catalyze the reduction of di-eosin-oxidized glutathione (GSSG) in a continuous fluorescent reductase activity assay (with an apparent IC_50_ of 19.2 µM) [28]. Others have reported apparent IC_50_ values for punicalagin in the di-eosin-GSSG assay at 6.1 µM [35]. To further compare the inhibitory effect of punicalagin and to assess its substrate specificity, we performed an insulin reduction assay. This assay evaluates PDI’s capacity to cleave disulfide bonds between insulin chains, which leads to increased solution turbidity over time [37]. Recombinant PDI was incubated with increasing concentrations of punicalagin (up to 600 µM), and absorbance changes were monitored. Inhibition increased with rising punicalagin concentrations, yielding an apparent IC₅₀ value of approximately 21 µM (**Figure 1B**), which is slightly higher than the reported IC_50_ value of 13.7 µM for insulin reduction [35]. This result supports punicalagin as a potent antagonist of PDI reductase activity and indicates that its effect is not restricted to a specific substrate, such as GSSG in the fluorescence reductase activity assay [28].

PDI’s reductase activity is linked to its ability to transfer disulfides via its oxidase activity and the effect of punicalagin on PDI’s oxidase activity has never been investigated. To test whether punicalagin also affects PDI oxidase activity, we employed a decapeptide oxidation assay using the synthetic peptide NRCSQGSCWN. Disulfide bond formation within this peptide quenches intrinsic Trp fluorescence via proximity to the positively charged Arg, allowing continuous measurement of PDI’s oxidase activity over time [38, 39]. In this experiment, recombinant PDI was incubated with punicalagin under redox-buffered conditions (GSH:GSSG of 4:1) prior to initiating oxidase activity by adding the peptide substrate. PDI in the presence of vehicle control produced the greatest fluorescence decay, whereas the addition of punicalagin led to a rate similar to peptide only-no enzyme control conditions, indicating substantial inhibition of PDI’s oxidase activity by punicalagin (**Figure 1C-D**).

As PDI’s oxidoreductase activity are dependent on the active site cysteines within its CGHC motifs, we evaluated whether punicalagin directly modifies the redox state of PDI’s catalytic cysteines. We performed free thiol labeling with a maleimide–PEG2–biotin reagent. Maleimide is an electrophilic warhead that targets the nucleophilic sulfur atom of the thiol of reduced cysteines[40]; the biotin functionalization of maleimide allows for detection of the probe. First, recombinant PDI was first either pre-oxidized with the physiologic oxidant H₂O₂, or pre-reduced with phosphines (TCEP) prior to incubating it with increasing concentrations of punicalagin. In pre-oxidized PDI, punicalagin did not increase maleimide labeling relative to vehicle control, indicating that punicalagin does not chemically reduce disulfide bonds formed on the CGHC cysteines. Similarly, in pre-reduced PDI, punicalagin failed to decrease thiol labeling, showing that the compound does not oxidize the catalytic cysteines even at high concentrations (**Figure 1E**). Therefore, punicalagin does not directly affect the intrinsic redox chemistry of PDI’s active sites.

Taken together, these findings demonstrate that punicalagin inhibits both reductase and oxidase activities of PDI with micromolar potency but does so without directly modifying or altering the redox state of the enzyme’s catalytic thiols. The results support a non-covalent or allosteric mechanism of inhibition and establish punicalagin as a natural modulator of PDI’s catalytic function, providing a mechanistic framework for additional studies on galloylated polyphenols and redox enzyme regulation.

### 3.2. Molecular docking predicts punicalagin binding sites on PDI

We and others have found that punicalagin antagonizes several members of the thiol isomerase family members [28, 34, 35]. To investigate the molecular basis of PDI inhibition by punicalagin, we performed unbiased molecular docking simulations using the full-length crystal structure of PDI (PDB: 4EKZ). Blind docking identified multiple candidate binding sites across the protein, visualized as regions R1 to R9 in **Figure 2A** The top-scoring poses indicated that punicalagin preferentially interacts with the N-terminal **a** domain (binding energy: –9.4 kcal/mol; R1) and (–9.0 kcal/mol; R2), with weaker predicted binding to rest of the protein (between -8.8 and –7.9 kcal/mol; R3-R9) (**Table 1**). Binding to the other **b, b’,** and **a’** domain were also apparent and were consistent with previous study observing that punicalagin binds to both the **a** and **a’** domains [34]. Detailed analysis revealed that punicalagin forms diverse molecular interactions with PDI in the N-terminal **a** domain, including hydrogen bonds, salt bridges, and hydrophobic contacts. Among these, hydrogen bonds were the predominant interaction type (**Figure 2B**). The spatial clustering and energetic ranking of the binding sites suggest that punicalagin can engage multiple surface regions in the N-terminal domain. Overall, these results provide a structural rationale for the inhibitory effects of punicalagin on PDI activity and highlight alternative regions and interaction types that may be important for future structure-based design of inhibitors.

**Figure 2.**
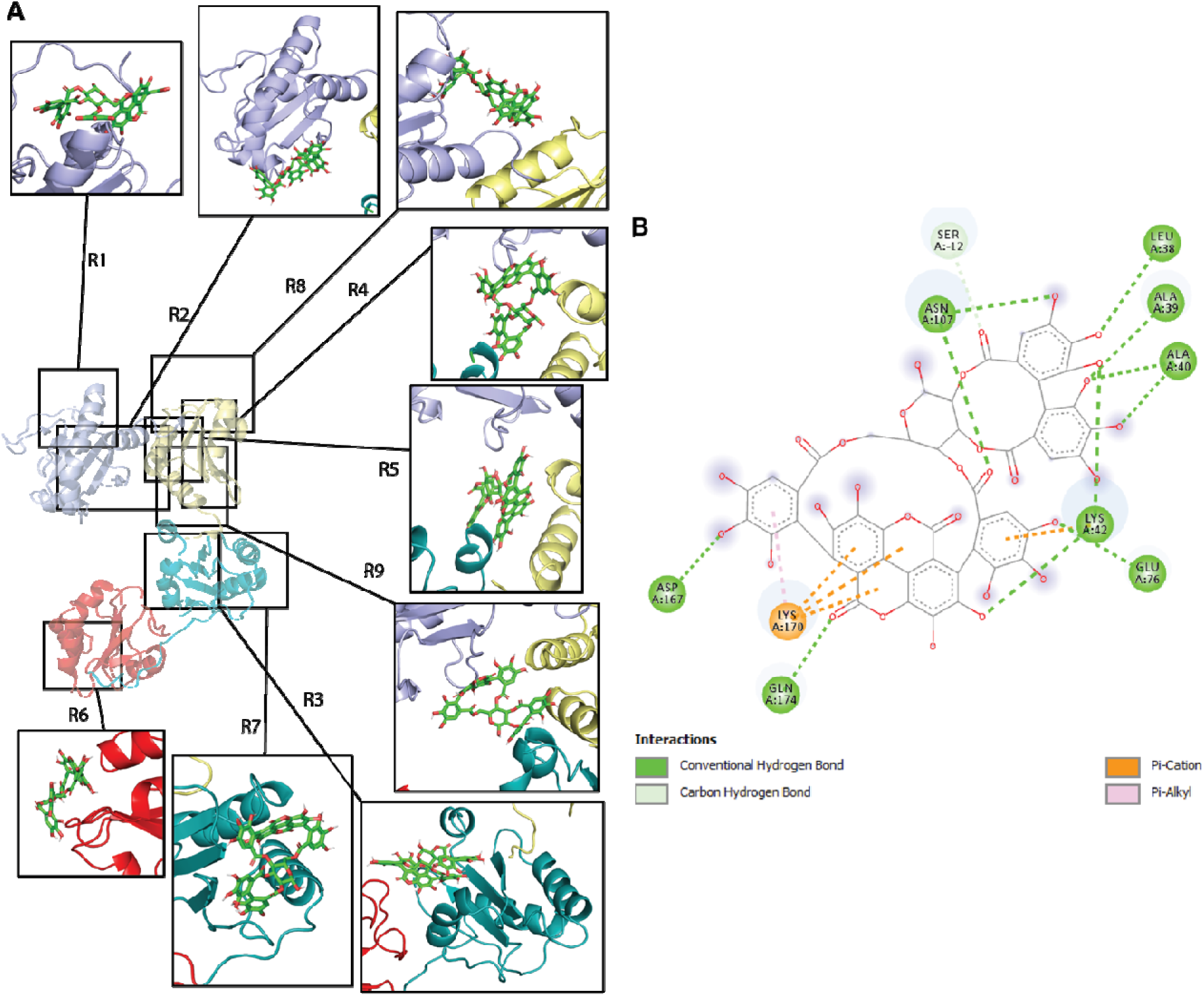
Punicalagin’s predicted binding to PDI by molecular docking. (A) Nine different binding poses are thermodynamically favorable. Many of the interactions occur within the vicinity of the a and b domains. Grey, a domain; yellow, b domain; green, b’ domain; red, a’ domain. (B) Molecular Interactions associated with R1. R = region.

### 3.3. Molecular dynamics simulation reveals that punicalagin binds stably to the **a** domain of PDI

To further elucidate the dynamic features of the PDI–punicalagin complex, we performed all-atom molecular dynamics (MD) simulations in explicit water at 37°C, 1 bar pressure, and pH 7.2. Interaction snapshots were collected from 0 ns to 500 ns, as illustrated in **Figure 3A** and in the **Supplementary Video**. Throughout the simulation, punicalagin consistently engaged residues located predominantly in the **a** domain with intermittent interactions extending into the **b** domain. This interaction supported structural shift to PDI by adopting an open configuration via extending the **a’** domain.

**Figure 3.**
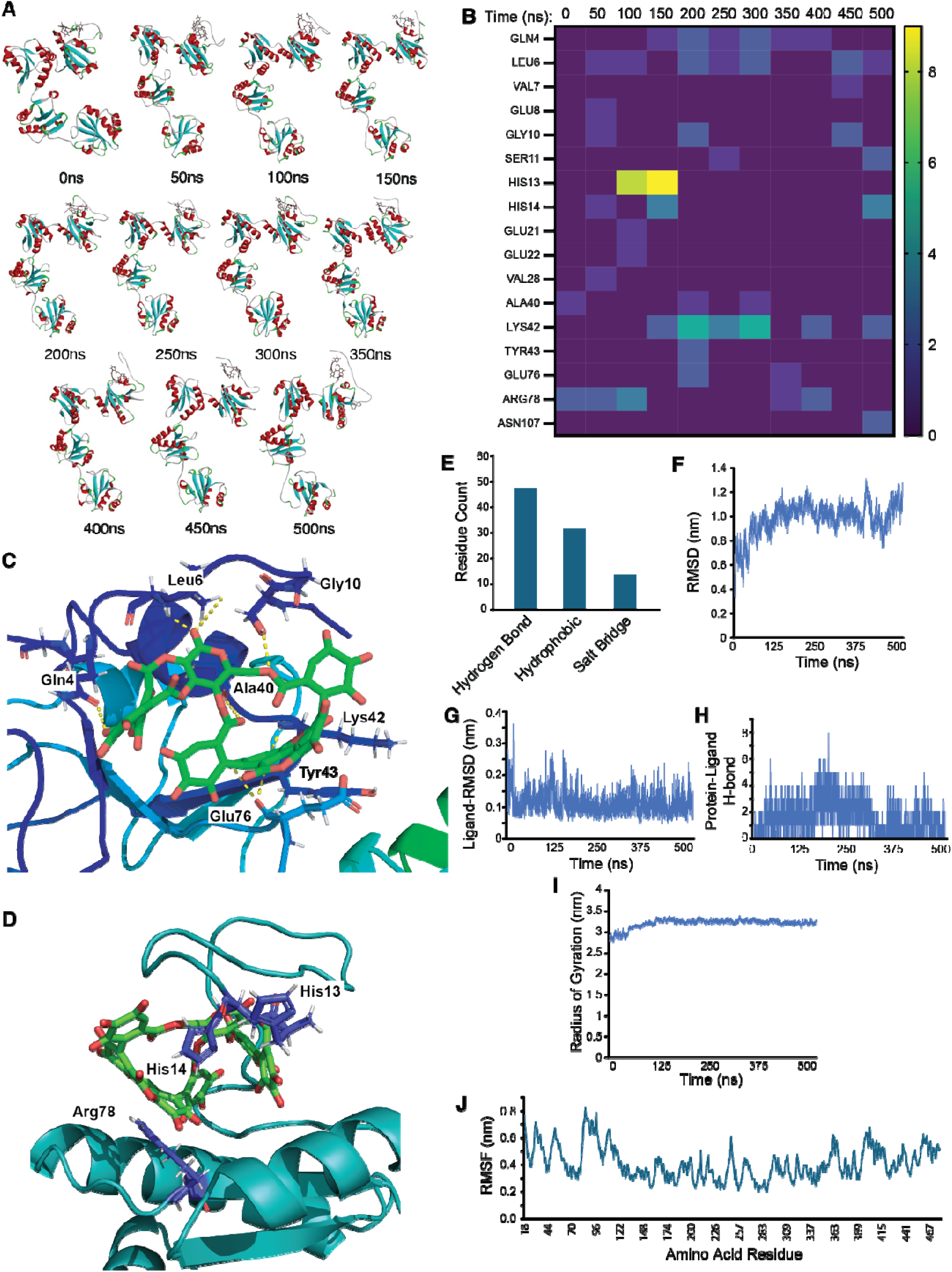
Molecular dynamics simulation reveals stable binding of punicalagin to the a domain of PDI predominantly through hydrogen bond interaction. (A) 50 ns kinetic time frames of punicalagin binding to PDI (PDB: 4ekz) during molecular dynamic simulation. (B) Heat map of the molecular dynamic simulation showing the number of residue contact over time. (C) Molecular interactions of punicalagin to PDI residues at 200 ns. (D) Molecular interactions of punicalagin to PDI His13, His 14, and Arg78 residues at 150 ns. (E) Distribution of the molecular interaction types onto PDI. (F) Protein root-mean-square deviation (RMSD) during the simulation. (G) Punicalagin RMSD during the simulation. (H) H-bond interactions between PDI and punicalagin throughout the simulation. (I) Radius of gyration of the interaction. (J) Root-mean-square-fluctuation of the amino acid residues during the simulation.

To quantify interaction frequency and specificity, we constructed a heat map (**Figure 3B**) depicting residue contacts over time. The periods of maximal interaction were observed at 100 ns and 150 ns, as reflected by the highest number of contact occurrences. Analysis at 100 and 150 ns revealed stable interactions with His13, and to a lesser extent, with His14 and Arg78 (**Figure 3B**). By 200 ns, punicalagin engaged an expanded set of residues, notably Leu6, Gln4, Ala40, Glu76, Tyr43, Lys42, and Gly10, primarily though hydrogen bonding. Further inspection indicated that the ellagic acid group of punicalagin formed hydrogen bonds with Lys42 (backbone amide to ellagic acid hydroxyl oxygen), Tyr43 (backbone amide to hydroxyl oxygen), and Glu76 (backbone carboxyl oxygen to ellagic acid hydroxyl hydrogen) (**Figure 3C**). Meanwhile, the gallic acid moieties preferentially interacted with Leu6, Gln4, and Ala40 via backbone amide or carboxyl group hydrogen bonds, as well as with Ser11 and Gly10 through sidechain and amide group contacts (**Figure 3C**). His13, His14, and Arg78 interacted with punicalagin through hydrogen bonding and Pi-stacking interactions (**Figure 3D**). Collectively, these results underscore the critical anchoring role of multiple hydrogen bonds, particularly those involving polyphenol hydroxyl groups, in punicalagin’s binding to PDI. Diverse noncovalent interactions, including salt bridges and hydrophobic contacts, were also detected, but hydrogen bonds predominated overall (**Figure 3E**). Notably, these hydrogen bonding events were spatially clustered to the **a** domain, supporting its principal role in punicalagin recognition.

Trajectory analysis further verified the stability of the complex. The protein root-mean-square deviation (RMSD), ligand RMSD, protein–ligand hydrogen bond counts, and radius of gyration (**Figure 3F-I**) indicated relatively stable binding, with the highest root-mean-square deviation (RMSD) reaching 1.22 nm at ∼400 ns and ∼500 ns, coinciding with conformational rearrangements in the complex. Most RMSD values remained less than 1.4 nm, reflecting interim stability and a lack of major structural disruption in the ligand–protein interface. Additionally, root-mean-square fluctuation (RMSF) analysis revealed that the interaction with punicalagin did not induce excessive flexibility in any single region of PDI, reinforcing the notion of a defined and stable binding site (**Figure 3J**).

Energetic analysis using the molecular mechanics Poisson–Boltzmann surface area (MM–PBSA) method predicted a favorable binding free energy (ΔG_total) for the PDI–punicalagin complex of –19.16 ± 0.85 kcal/mol (**Table 2**). The major contributions to binding energy arose from van der Waals (ΔE_vdw = –42.38 ± 0.47 kcal/mol) and electrostatic interactions (ΔE_eel = –16.27 ± 0.49 kcal/mol). The polar solvation contribution (ΔE_PB = 44.35 ± 0.94 kcal/mol) was largely offset by favorable nonpolar solvation effects (E_npolar = –4.86 ± 0.15 kcal/mol), consistent with a binding mode stabilized by numerous hydrogen bonds and complementary surface interactions.

Taken together, these MD simulations provide compelling evidence that punicalagin binds stably to PDI’s **a** domain, predominantly bound by hydrogen bonding and van der Waals contacts involving both its ellagic acid and galloyl groups.

### 3.4. Network analysis (Plex) reveals PDI and broader signaling targets of galloylated polyphenols

To systematically explore molecular targets of punicalagin (PC), we employed Plex, a transparent artificial intelligence (AI) platform powered by a comprehensive biomedical knowledge graph, spanning nearly one billion nodes and edges across pharmacology, chemical biology, and multi-omics datasets [36]. Plex uses focal graphs to rapidly synthesize information from diverse sources, generating high-confidence hypotheses based on the convergence of independent evidence. Using the Plex AI platform, we constructed a focal graph based on chemical structure similarity analysis for PC (**Figure 4A**). Pinocembrin 7-O-(3”-galloyl-4”,6”-(S)-hexahydroxydiphenoyl)-β-D-glucose (PGHG) was also used as a query control compound since we previously published PGHG as a galloylated polyphenol antagonist of thiol isomerases [28]. This approach integrates molecular information from large-scale biomedical knowledge to highlight relationships between compounds and its potential protein targets. We further generated a sub-focal graph to visualize specific connections among PC and PGHG (shown in yellow), similar chemical structures (shown in red), and predicted protein targets (shown in blue) (**Figure 4B**). Analysis using this platform revealed PDI (gene name prolyl 4-hydroxylase beta chain; P4HB), as a primary predicted target for both PC and PGHG (**Figure 4B**). This outcome strengthens and independently validates our experimental identification of PDI as a direct target for PC. Other predicted targets included various isoforms of protein kinase C (PKC) and members of the mitogen-activated protein kinase (MAPK) family. These findings were visualized in the sub-focal graph (**Figure 4B**), highlighting a broader molecular target profile for galloylated polyphenols. Experimental data confirmed that both PC and PGHG directly inhibit protein PDIs and effectively reduce platelet aggregation induced by agonists that engage PKC and MAPK signaling pathways [28, 41, 42]. Our Plex focal graph analysis indicates that galloylated polyphenols such as PC and PGHG target multiple proteins. This focal graph approach, efficiently and concisely integrating data from multiple large-scale biomedical databases, supports our experimental findings and provides a comprehensive network of compound–target relationships.

**Figure 4.**
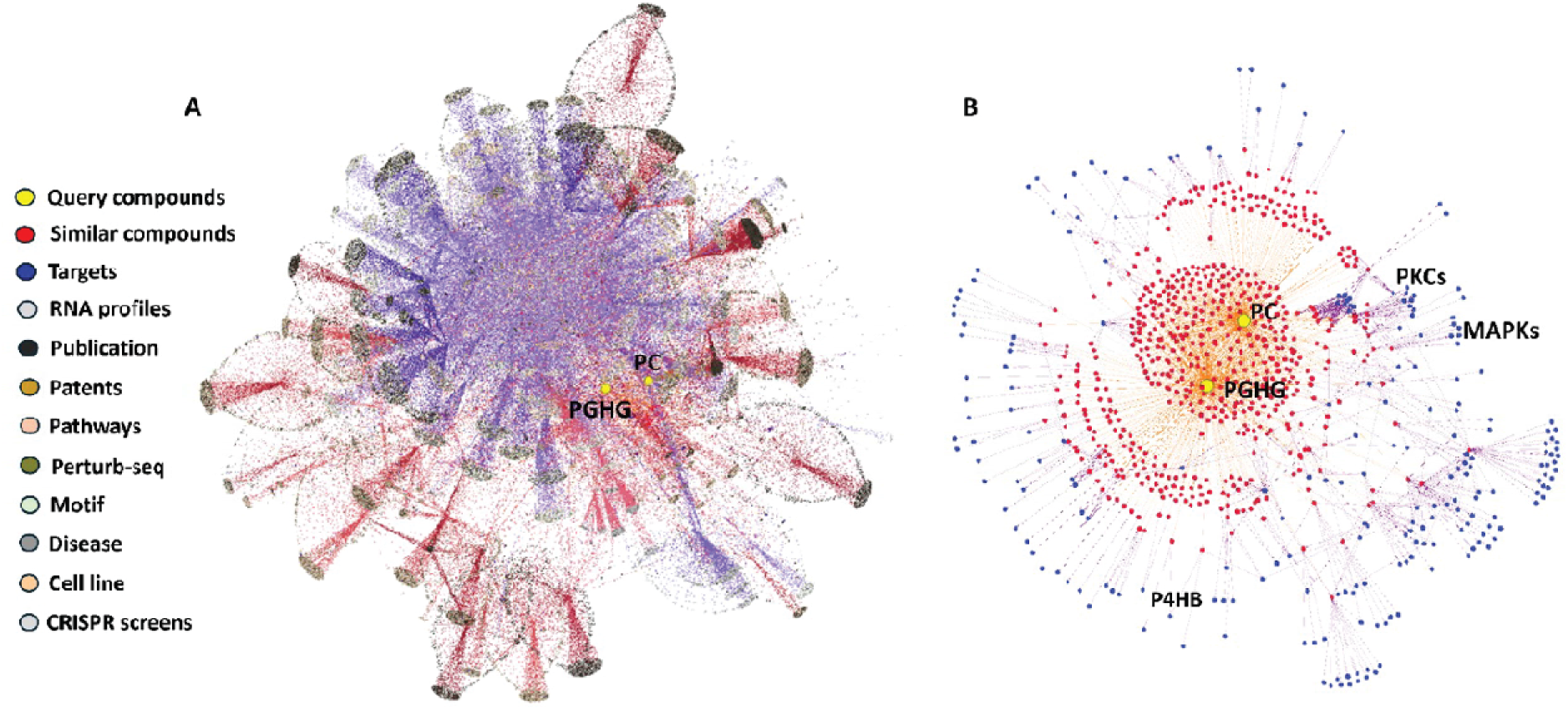
Focal graph analysis of punicalagin (PC) and the pinocembrin-based galloylated polyphenol (PGHG). (A) Focal graphs of Punicalagin (PC) and PGHG as the query compounds showing connections to diverse data types (indicated by the color legend). (B) A subgraph of the focal graph is shown on the right comprising the connections between only the query compounds (Yellow, PC and PGHG), similar compounds (red), and targets (blue). Representative top ranked targets are labeled. Focal graph images were generated by Gephi (https://gephi.org).

## 4. Discussion

Punicalagin is a structurally complex galloylated polyphenol that has been linked to antithrombotic, anti-inflammatory, and cytoprotective effects in multiple preclinical models [28–31]. PDIs are central redox hubs at the intersection of protein folding, thrombosis, and inflammatory signaling, functioning both in the endoplasmic reticulum and at the cell surface to regulate disulfide exchange in client proteins, integrins, and platelet receptors [13–15, 17, 18, 43, 44]. The present study supports a model in which punicalagin functions as a noncovalent, domain-selective inhibitor of PDI oxidoreductase activity, acting through stable binding to N-terminal domains and modulation of conformational dynamics rather than through direct modification of catalytic cysteines.

Prior work showed that punicalagin binds PDI and ERp57 with similar affinity and inhibits their disulfide reductase activity, with greater efficacy toward ERp57 [34, 35]. In these studies, docking and molecular dynamics simulation were performed to propose binding pockets in the redox **a**′ domain and to relate ligand binding to changes in protein thermal stability [34]. More recently, this framework was extended by comparing punicalagin with the structurally simplified ellagitannin punicalin that does not contain the hexahydroxydiphenic acid moiety. This study reported that punicalin retains ERp57 binding and inhibition but exhibits markedly reduced activity toward PDI, thereby sharpening isoform selectivity and further implicating redox domains as key determinants of ligand recognition[35]. Together, these studies suggest punicalagin and related ellagitannins as noncovalent PDI inhibitors and provide important insight into isoform preference and likely binding regions, particularly within redox-active domains. Building on this foundation, this present work focuses on complementary questions, using long-timescale molecular dynamics, oxidase assay, thiol-resolving biochemical assays, and AI-guided network analysis to characterize catalytic cysteine redox state during inhibition, effects on oxidase activity, domain-resolved conformational ensembles, and the positioning of PDI within broader punicalagin-responsive signaling networks relevant to thrombo-inflammatory pathways.

In this study, punicalagin inhibited oxidoreductase activities across distinct substrates, including glutathione-based probes [28], insulin, and the synthetic decapeptide NRCSQGSCWN, without altering active-site thiol labeling. This indicates that its effects are not substrate restricted. This study probes the effect of punicalagin on the oxidase function of PDI, which has not been previously elucidated because of the technical challenges associated with the oxidase assay [27]. In this context, the oxidase assay was also inhibited, indicating a dual inhibitory effect of punicalagin on both reductase and oxidase activities. The preserved catalytic redox state observed via thiol labeling indicates that punicalagin does not promote net oxidation, reduction, or covalent masking of CXXC motifs, arguing against simple redox trapping or electrophilic adduction as primary mechanisms. The prior studies did not directly quantify catalytic thiol redox state under inhibitory conditions, and our structural and dynamic analyses provide mechanistic support for this model [34, 35]. Docking and molecular dynamics clustered punicalagin binding to the **a** and **b** domains, with energetically favorable poses towards the **a** domain and sustained contacts involving residues such as Leu6, Gln4, His13, Tyr43, Lys42, Glu76, Ser11, and Gly10. The ellagic and galloyl groups formed dense hydrogen-bond networks and van der Waals contacts along these surfaces, consistent with the high hydroxyl density and aromatic character that define galloylated polyphenols [29, 30]. The trajectories also revealed periods of increased backbone deviation associated with more open domain arrangements, in line with studies demonstrating that PDI samples compact and open conformations that are sensitive to redox state and ligand binding [45–47]. In this context, punicalagin appears to stabilize an alternative (open) conformational ensemble that is compatible with binding but less efficient for catalysis, thereby weakening oxidoreductase throughput without chemically “locking” the active site through covalent interaction. In contrast to previous studies that predominantly localized punicalagin binding to redox active C-terminal domains (**a’** domain), our docking and extensive molecular dynamics simulations clustered punicalagin at the N terminal **a** and **b** domains, with particularly favorable poses within the **a** domain. In this context, we provide evidence on alternative binding modalities that may be important to characterize the mechanism of inhibition.

Covalent inhibitors such as PACMA31, RB-11-ca and CCF642 which act as irreversible small-molecule inhibitors of PDI by forming a covalent bond with active-site cysteines, exemplify an alternative mode of PDI inhibition in which the CXXC motif is modified [7, 48–50]. Alternatively, the preservation of catalytic thiol redox state in the presence of punicalagin aligns it with a growing class of noncovalent PDI inhibitors that act through domain-specific or allosteric mechanisms [15, 45, 46, 51–53]. For example, bepristat 2a and LOC14 binds outside the catalytic CXXC motifs and modulates PDI activity through changes in conformational dynamics and allosteric coupling [45, 51, 52], while isoquercetin and related flavonoids also inhibit extracellular PDI, reduce thrombin generation, and attenuate venous thrombosis *in vivo* without modifying catalytic cysteines [15, 46, 53, 54]. The combination of substrate-independent inhibition, molecular docking, molecular dynamics simulations, and unchanged thiol redox chemistry supports a model in which punicalagin interferes with conformational dynamics, substrate engagement, or intradomain communication rather than directly blocking the redox chemistry of the active site. In doing so, this work extends the paradigm of noncovalent, allosteric PDI inhibition to a large, dietary ellagitannin and provides residue-level hypotheses for how such compounds differentially engage PDI domains.

The Plex-based network analysis situates punicalagin and another galloylated polyphenol, PGHG, within a broader signaling framework. P4HB (canonical PDI) emerged as a convergent, high-confidence node for both compounds, independently validating PDI as a core target, while protein kinase C isoforms and mitogen-activated protein kinases were also highlighted as candidate targets. These predictions align with experimental data showing that galloylated polyphenols and other tannins suppress PKC- and MAPK-dependent inflammatory and platelet signaling, including inhibition of platelet activation and aggregation by ellagitannins and catechin gallates [20, 28–30]. From this perspective, PDI inhibition appears to be part of a coordinated modulation of thiol isomerases and downstream kinase cascades that collectively recalibrate thrombotic and inflammatory responses rather than a single-enzyme effect.

Phytochemical and pharmacological studies have established punicalagin as a major ellagitannin constituent of pomegranate peels, Indian almond leaf extracts, and velvet bushwillow bark with antioxidative, anti-inflammatory, antithrombotic, and antitumor activities in diverse cells and animal models [28–30]. These studies demonstrate favorable effects on vascular function, inflammation, and cancer-relevant pathways and generally report low toxicity at doses achievable through dietary or formulated administration. However, despite growing interest in thiol isomerases as druggable targets, the pharmacology of punicalagin specifically in the context of PDI inhibition remains incompletely defined. Existing work has focused largely on global outcomes (for example, reduced oxidative stress, platelet activation, or tumor growth) without dissecting how parent punicalagin versus its metabolites (ellagic acid, urolithins) distribute *in vivo* reach thiol isomerases, or differentially modulate PDI activity in tissues. We previously found that punicalagin administered by oral gavage in mice prevented thrombus formation following chemical injury to the carotid arteries [28]. Specifically, administration of the compound at 100 mg/kg once by gavage increased the compound concentration in the plasma to approximately 100 ng/mL by 60 and 90 min post-gavage [28]. Despite this information, the dose–exposure–response relationship, long-term safety, and isoform selectivity of punicalagin as a thiol isomerase modulator are still unclear and will require integrated pharmacokinetic, metabolite profiling, and target-engagement studies to resolve. In addition, a case report suggested punicalagin engages with therapeutic doses of the anticoagulant warfarin potentially through the cytochrome P450 system [55]. Such engagement underscores the importance to further understand the antithrombotic mechanism of the compound *in vivo*.

Overall, this study establishes punicalagin as a micromolar, noncovalent antagonist of PDI oxidoreductase activity that acts through defined N-terminal binding sites and conformational modulation, rather than catalytic cysteine redox modification. By combining structural, biochemical, and network-level analyses, the work advances a cohesive mechanistic picture of how galloylated polyphenols can shape thiol isomerase function and associated signaling pathways. In addition, this work strengthens the rationale for leveraging galloylated polyphenols as foundational scaffolds in the development of redox-centered therapeutics for thrombosis, inflammatory, and proteostasis-related diseases.

## CRediT authorship contribution statement

Conceptualization, O.C.O., M.Y.; Data Curation, O.C.O., O.L., M.Y.; Formal Analysis, O.C.O., O.L., M.Y.; Funding Acquisition, O.L., P.D.-C., M.Y.; Investigation, O.C.O., Q.P.K., I.H.Z., O.L.; Methodology, O.C.O., O.L.; Project Administration, M.Y.; Resources, O.L., P.D.-C., M.Y.; Supervision, M.Y.; Validation, O.C.O., M.Y.; ; Visualization, O.C.O., O.L., M.Y.; Writing – Original Draft, O.C.O., Q.P.K., O.L., P.D.-C., M.Y.; Writing – Review & Editing, O.C.O., O.L., P.D.-C., M.Y.

## Consent for publication

All the authors have read and approved the manuscript for publication.

## Declaration of competing interest

M.Y. has a patent pending with Robert Flaumenhaft at Beth Israel Deaconess Medical Center entitled, “Compounds and Methods for Using Galloylated Polyphenols to Treat Diseases Mediated by Thiol Isomerases”. O.L. is a part-time paid consultant to Plex Research; the other authors declare no relevant conflict of interest.

## Declaration of generative AI and AI-assisted technologies

During the preparation of this work, the authors used Plex AI to understand the interaction of galloylated polyphenols with different molecular pathways. After using this tool or service, the authors reviewed and edited the content as needed and take full responsibility for the content of the publication.

## Acknowledgements

This work was supported by the National Institute of Health grants R00HL164888 (M.Y.), R00HL177831 (P.D.-C.), a Zoll Foundation Award (O.L.), and a Bloodworks Northwest investigator startup fund (M.Y.). These funding sources had no involvement in writing this article or submitting this article for publication.

## Appendix A. Supplementary data

The following is the supplementary data to this article:

Multimedia component 1.

## Data availability statement

The data presented in this study are available on request from the corresponding author.

## Abbreviations

PDI: protein disulfide isomerase
ER: endoplasmic reticulum
SBP: streptavidin binding protein
TB: terrific broth
IPTG: isopropyl β-D-1-thiogalactopyranoside
PBS: phosphate-buffered saline
TFA: trifluoroacetic acid
GSH: reduced glutathione
GSSG: oxidized glutathione
DTT: dithiothreitol
RMSD: root-mean-square deviation
RMSF: root-mean-square fluctuation
AI: artificial intelligence
PC: punicalagin
PGHG: Pinocembrin 7-O-(3”-galloyl-4”,6”-(S)-hexahydroxydiphenoyl)-β-D-glucose
MD: molecular dynamics
MM-PBSA: molecular mechanics Poisson-Boltzmann surface area
TCEP: phosphine
PKC: protein kinase C
MAPK: mitogen-activated protein kinase.

