## Supplemental Materials for "Molecular insights into protein disulfide isomerase antagonism by punicalagin"

**Supplemental Contents**

Supplemental Materials . . . . . . . . . . . . . . . . . . . . . . . . . . . . .. . . . . . . 2

Supplemental Video . . . . . . . . . . . . . . . . . . . . . . . . . . . . .. . . . . . . . . 4

**Supplemental Materials**

**Reagents**

PDI monoclonal antibody (RL90) (Cat. No. MA3-019), EDTA-free (100x) Halt Protease/Phosphatase inhibitor (Cat. No. 78443), Pierce High-Capacity Streptavidin Agarose (Cat. No. 20361), Trifluoroacetic acid (TFA, Cat. No. 28901), EZ-Link™ Maleimide-PEG2-Biotin, No-Weigh™ Format (Cat. No. A39261), Membrane filters (Cat. No. DS0210-4020), and SuperSignal WestPico PLUS Chemiluminescent Substrate (Cat. No. 34580) were from ThermoFisher Scientific. Peroxidase AffiniPure™ Donkey Anti-Mouse IgG (H+L) (Cat. No. 715-035-150) was from Jackson ImmunoResearch Laboratories. Streptavidin-HRP conjugate (Cat. No. DY998) was from Bio-Techne. BL21(DE3) Competent Cells (Cat. No. C2527H), NdeI (Cat. No. R0111), and BamHI (Cat. No. R0136) were from New England Biolabs. Isopropyl β-D-thiogalactopyranoside (IPTG, Cat. No. I2481C50) was from Gold Biotechnology. Dithiolthreitol (DTT, Cat. No. 1610611), 4X Laemmli Sample Buffer (Cat. No. 1610747), Precast Gel, 4-15%MP TGX, 10W (Cat. No. 4561084DC), and Nitrocellulose membranes (Cat. No. 1704271) were from BioRad. Trizma hydrochloride (Tris-HCl, Cat. No. T3253-1KG), Sodium chloride (NaCl, Cat. No. S9888-5KG), Potassium Phosphate Monobasic (KH₂PO₄, Cat. No. P5655-500G), Potassium Phosphate Dibasic (K₂HPO₄, Cat. No. S3264-250G), Terrific broth (Cat. No. T9179-1KG), L-Glutathione reduced (GSH, Cat. No. G6013-10G), L-Glutathione oxidized (GSSG, Cat. No. G4376-5G), Acetonitrile (Cat. No. 34851-4L), Punicalagin (Cat. No. PHL80524-10MG), Hydrogen peroxide (H₂O₂, Cat. No. 88597-100ML-F), Tris(2-carboxyethyl)phosphine (TCEP, Cat. No. C4706-10G), and pT7-FLAG-SBP1 (Cat. No. P3871) was from Sigma Aldrich. Ethylenediaminetetraacetic acid (EDTA, Cat. No. MT-46034Cl) and Phosphate-Buffered Saline (PBS, Cat. No. 10010-023) were from Fisher Scientific. Decapeptide substrate (NRCSQGSCWN, Cat. No. Custom peptide synthesis) was from GenScript. Pinocembrin 7-O-(3''-galloyl-4'',6''-(S)-hexahydroxydiphenoyl)-β-D-glucose (PGHG, Cat. No. CFN90881) was from ChemFaces. Graphpad Prism v10 (<https://www.graphpad.com/features>) was from Graphpad by Dotmatics. PyMol (<https://pymol.org/2/>) was from Schrodinger. Autodock Vina (<https://vina.scripps.edu/>) was from Scripps University. GROMACs version 2020.6 (GPL versions) was from Uppsala University, University of Groningen. Discovery Studio Visualizer (<https://www.3ds.com/products/biovia/discovery-studio/visualization>) was from Dassault Systèmes BIOVIA. Gephi (<https://gephi.org>) was from Open source (Gephi Consortium). Plex AI (<https://www.plexresearch.com>) was from Plex Research Inc.

**Supplemental Video**

**Supplemental Video** – 500 ns molecular dynamics simulation of punicalagin binding to PDI (PDB: 4ekz). Stable binding of punicalagin to the N-terminal domain induces an open structural configuration on the enzyme.
